# *Drosophila* orthologues of oculocutaneous albinism-associated genes regulate sleep and circadian rhythm via visual neurotransmission

**DOI:** 10.64898/2026.05.27.728351

**Authors:** Mehran Akhtar, Yu-Chien Hung, Clifton Lewis, Nawel Medjadi, Johan Bence, Flaviano Giorgini, Mervyn G Thomas, Ko-Fan Chen

## Abstract

Melanin is a pigment found in the skin and cuticle of animals. Oculocutaneous albinism (OCA) is a group of autosomal recessive disorders defined by reduced melanin in skin and eyes, and is associated with visual defects such as foveal hypoplasia and infantile nystagmus. Sleep disturbance has been documented in children with OCA, and *oca2* loss-of-function in cavefish causes constitutive sleep loss, indicating a sleep regulatory function of OCA-associated genes in the visual system. To test potential roles of OCA-associated genes in regulating sleep and vision through evolutionarily conserved mechanisms, we used *Drosophila melanogaster*, a high-throughput phenotyping system to screen sleep and visual phenotypes for genetic mutants of OCA-associated genes. Among the OCA-associated genes, bidirectional DRSC Integrative Ortholog Prediction Tool identified Drosophila orthologues for *OCA2*, *SLC45A2*, *SLC24A5* and *LRMDA*. RNAi-mediated knockdown in developing *Drosophila* eye tissue identified the *OCA2* orthologues *hoe1*, *hoe2*, and the *SLC45A2* orthologue *lovit* as candidate genes required for normal sleep. Moreover, hoe1, hoe2 and lovit1 null alleles reduced sleep and circadian rhythmicity, and showed altered photoreceptor neurotransmission. Consistently, the enhancer-trap reporter assay and scRNA dataset revealed that these OCA orthologues express in brain regions critical for sleep and vision regulation. Collectively the data indicate evolutionarily conserved neuronal function of *OCA2* and *SLC45A2* orthologues that regulate sleep and photoreceptor neurotransmission.

**Significance Statement:** Oculocutaneous albinism (OCA) is a pigmentation disorder of the skin and eyes accompanied by reduced visual acuity. Children with albinism also report sleep disturbance, and the mechanism is unclear. We tested whether genes mutated in albinism regulate sleep in the fruit fly *Drosophila melanogaster*, an organism that has a divergent melanin synthesis pathway and lacks tyrosinase, the principal pigmentation enzyme in mammals. We found *Drosophila* OCA orthologues, *hoe1*, *hoe2* and *lovit* express widely in the nervous system and control sleep, circadian rhythmicity and neural activity of photoreceptors. Importantly no pigmentation defects were detected in aforementioned mutants, our finding is consistent with the proposal that pleiotropic neuronal function of OCA2 and SLC45A2 underlies the sleep and vision symptoms in albinism.

## Introduction

Melanin is a pigment found in skin, cuticle, hair and eyes across the animal kingdom. Variation of melanin amount generates different skin colours and the pigmentation patterns in vertebrates and insects, which are crucial for camouflage and mate recognition. Oculocutaneous albinism (OCA) is a group of autosomal recessive disorders defined by reduced melanin in the skin, hair and eyes. OCA is also associated with visual developmental defects including foveal hypoplasia, optic nerve misrouting and infantile nystagmus (1–3). Eight OCA subtypes have been described to date (OCA1–8) and linked to genetic mutations disrupting melanin synthesis. The genes mutated in these subtypes are *TYR* (OCA1), *OCA2* (OCA2), *TYRP1* (OCA3), *SLC45A2* (OCA4), *SLC24A5* (OCA6), *LRMDA* (OCA7) and *DCT* (OCA8). OCA5 has been mapped to chromosome 4q24 in a single Pakistani family but the causative gene remains unidentified (4).

Melanin biosynthesis diverges between mammals and insects. In mammalian melanocytes, tyrosine is oxidised by the rate-limiting enzyme tyrosinase (TYR) enzyme, which oxidises L-tyrosine to dopaquinone within specialised organelles called melanosomes. Dopachrome tautomerase (DCT) and Tyrosinase-related protein 1 (TYRP1) then catalyse subsequent steps in the eumelanin pathway downstream of dopaquinone. The membrane transporters OCA2 and SLC45A2 regulate melanosomal pH, which is required for tyrosinase activity (5, 6). SLC24A5 is a sodium/calcium/potassium exchanger that maintains melanosomal calcium balance (7). LRMDA encodes a leucine-rich repeat protein required for melanocyte differentiation (8). Tyrosinase orthologues are absent in insects and melanin pigmentation in their cuticles depends on conversion of tyrosine to L-DOPA/dopamine (9), which is then oxidised by phenoloxidases of a different protein family (10). This pathway is nevertheless similar to the neuromelanin synthesis in the mammalian neurons (11).

Beyond their well-established role in pigmentation, recent evidence suggests OCA-associated genes may have unexpected functions in nervous system development and sleep regulation (12–14). In particular, sleep disturbance is a recurrent clinical feature of visual impairment in childhood. Ingram et al. (12) surveyed 72 children with visual impairment and reported that 89% had Childhood Sleep Habits Questionnaire scores above the clinical cut-off (greater than 41). Within this cohort, children with OCA shared with other visual impairment disorders the same general sleep alterations with high CSHQ scores (50.5±7.6)) but scored lower on daytime sleepiness. Sleep loss has also been documented in *Astyanax mexicanus* cavefish, where loss-of-function variants in *oca2* cause both albinism and constitutive sleep reduction (15). The extent of visual developmental defect in fovea and optic nerve seen in OCA patients (16) as well as neuronal expression pattern of some OCA-associated genes (17, 18), also suggest pleiotropic function of these genes beyond pigment synthesis.

The consistent observation of visual defect across OCA subtypes may also contribute to the sleep alterations seen in albinism patients, since light and visual signalling regulates sleep and circadian behaviour across animal phyla. In mammals, rod, cone and the intrinsically photosensitive retinal ganglion cells (ipRGC) mediate circadian photoentrainment as well as producing both sleep-promoting and arousal-promoting responses to light depending on wavelength (19, 20). Separately, the ipRGC–preoptic projection mediates the acute effect of light on sleep (21). Changes in visual load have also been associated with altered sleep structure in vertebrates (22, 23). Similarly in *Drosophila*, both visual and non-visual photoreceptors are required for circadian entrainment (24). Moreover, visual experience drives sleep need through HS/VS motion-processing neurons (25), and phototransduction promotes daytime sleep through R1–R6 eye photoreceptors and their downstream neural targets (26).

These findings raise the possibility that OCA-associated genes regulate sleep through evolutionarily conserved mechanisms in the eyes that are partly independent of melanin synthesis itself. To test this possibility, we used the established insect sleep model, *Drosophila melanogaster* (27), and screened for sleep phenotypes using RNAi-mediated knockdown in the developing eye. We then assessed knockout alleles for the candidates that emerged. We also recorded electroretinograms to test whether the observed sleep phenotype coincides with altered visual neurotransmission. We report that *OCA2* and *SLC45A2* orthologues regulate sleep and photoreceptor neurotransmission in a system that lacks the melanocyte lineage, indicating a pigmentation-independent neuronal function of these transporter genes.

## Results

### Bidirectional orthology mapping for OCA1–8 genes

Of the seven known OCA-associated genes screened, three (*TYR*, *TYRP1*, and *DCT*) had no *Drosophila* orthologue and could not be tested in this system. The remaining four OCA-associated genes (*OCA2*, *SLC45A2*, *SLC24A5*, *LRMDA*) returned bidirectional best-hit orthologues. *OCA2* mapped to two paralogues, *hoe1* and *hoe2*; both belonging to the CitM (TC 2.A.11) transporter superfamily, the same family as human OCA2 (5). *SLC45A2* mapped to *Slc45-1* and *lovit*. The other genes mapped to a single fly orthologue each: *LRMDA* → *CG31076*, *SLC24A5* → *CG12061*. To verify any roles in pigmentation for these OCA orthologues, we examined the publicly available phenotype database [https://bristlescreen.imba.oeaw.ac.at, (28)] where RNAi-based knockdown of candidate genes in developing cuticles was performed.(28) Only the knockdown of LRMDA caused mild darkening of pigmentation, consistent with the notion that the function of *Drosophila* OCA orthologues diverges from pigment synthesis. We therefore used the Gal4-UAS system to conduct RNAi-mediated knockdown for *Drosophila LRMDA*, *SLC24A5*, *OCA2* and *SLC45A2* orthologues and monitored their sleep profile and parameters (see Methods).

### RNAi-mediated knockdown of OCA orthologues

#### Knockdown of LRMDA and SLC24A5 orthologues does not cause a sleep phenotype

Targeting *LRMDA/CG31076* via three independent RNAi lines (TRiP, KK, GD) produced no changes in sleep parameters that met the two-control criterion in either sex (Figure S1; Table S2). Two RNAi line (GD, TriP) was tested for *SLC24A5/CG12061*, but again no sleep parameters showed significant changes in compared with controls (Figure S2; Table S2).

#### Knockdowns of either OCA2 orthologues (hoe1/2) cause inconsistent sleep phenotypes

Three RNAi lines (TRiP9, TRiP7, KK) targeting *hoe1* were tested. No line produced a sleep parameter change that was significant in comparison to controls in males (Figure S3A-C; Table S2). However, expression of TRiP9 RNAi caused significant day sleep loss in females (Figure S3CIIi).

Two RNAi lines (TRiP, KK) targeting *hoe2* were tested. The TRiP line reduced daytime sleep and daytime sleep bout length relative to both controls in males (Figure S4B). The KK line did not replicate this finding. The *hoe1/2* RNAi result is suggestive but not validated by independent line replication (Table S2).

#### Knockdowns of the SLC45A2 orthologue (lovit) causes sleep loss

Two RNAi lines (KK, GD) targeting *Slc45-1* produced no sleep parameter change against both controls in either sex (Figure S5; Table S2). Five independent RNAi lines (RNAi^1-5^) targeting *lovit* were tested. Expressing RNAi^1^ caused day sleep bout reduction in females (Figure S6AII), while expression of RNAi^2^ caused no sleep phenotypes (Figure S6B). The expression of the remaining three RNAi lines caused significant reductions in sleep amount and bout length in the night in comparison to both controls in both sexes (RNAi^3-5^, Figure S6C-E).

### Null and hypomorphic alleles of *hoe1/2 and lovit* show sleep and circadian alterations

Based on the above genetic screen, sleep loss can be observed in some, but not all RNAi knockdowns of *hoe1*, *hoe2* and *lovit* in the developing eyes (Table S2). To verify the observed sleep phenotypes independently, we obtained confirmed null or potential hypomorphic alleles for these three genes and tested the sleep profile for these lines. We also visually inspected the cuticles for all the five lines obtained and did not observe any cuticle colouration changes.

#### hoe1/2 and lovit mutant alleles reduce sleep amount and maintenance

For *hoe1*, the 3370 CRSPR-based deletion allele showed reduced daytime sleep in males and reduced sleep bout length in day and night in both sexes (Figure 1B). The 62601 allele showed nighttime sleep loss and bout length reduction in both sexes, whereas only females showed sleep loss and bout length reduction in the day (Figure 1B). For *hoe2*, 2795 alleles showed reduced daytime sleep only in males and reduced sleep bout length in day and night in both sexes (Figure 1C). Males of the 92669 allele showed reduced day sleep whereas both males and females showed increased nighttime sleep. The sleep bout length of the 92669 allele is reduced in daytime in both sexes and in nighttime only in males (Figure 1C). Overall, *hoe1* and *hoe2* mutants exhibit poor sleep quality with either reduced bout length or total amount of sleep during the day. Sleep was measured in *lovit*^1^ males and females in the *iso31* background with *lovit*^1^ males and females showing reduced nighttime sleep (p < 0.0001) and sleep bout length (p < 0.05) as compared to *iso31* (Figure 1A Iii, Iiv, IIii, IIiv). While a clear lower peak sleep level at mid-day was observed in female *lovit^1^* flies (Figure 1AII), only male *lovit^1^* flies showed reduced daytime sleep (p < 0.01) and day sleep bout length (p < 0.05) (Figure 1AI-i, iii). Sleep parameters for *hoe1/2* and *lovit^1^*mutant alleles are summarised in Table S3.

**Figure 1.**
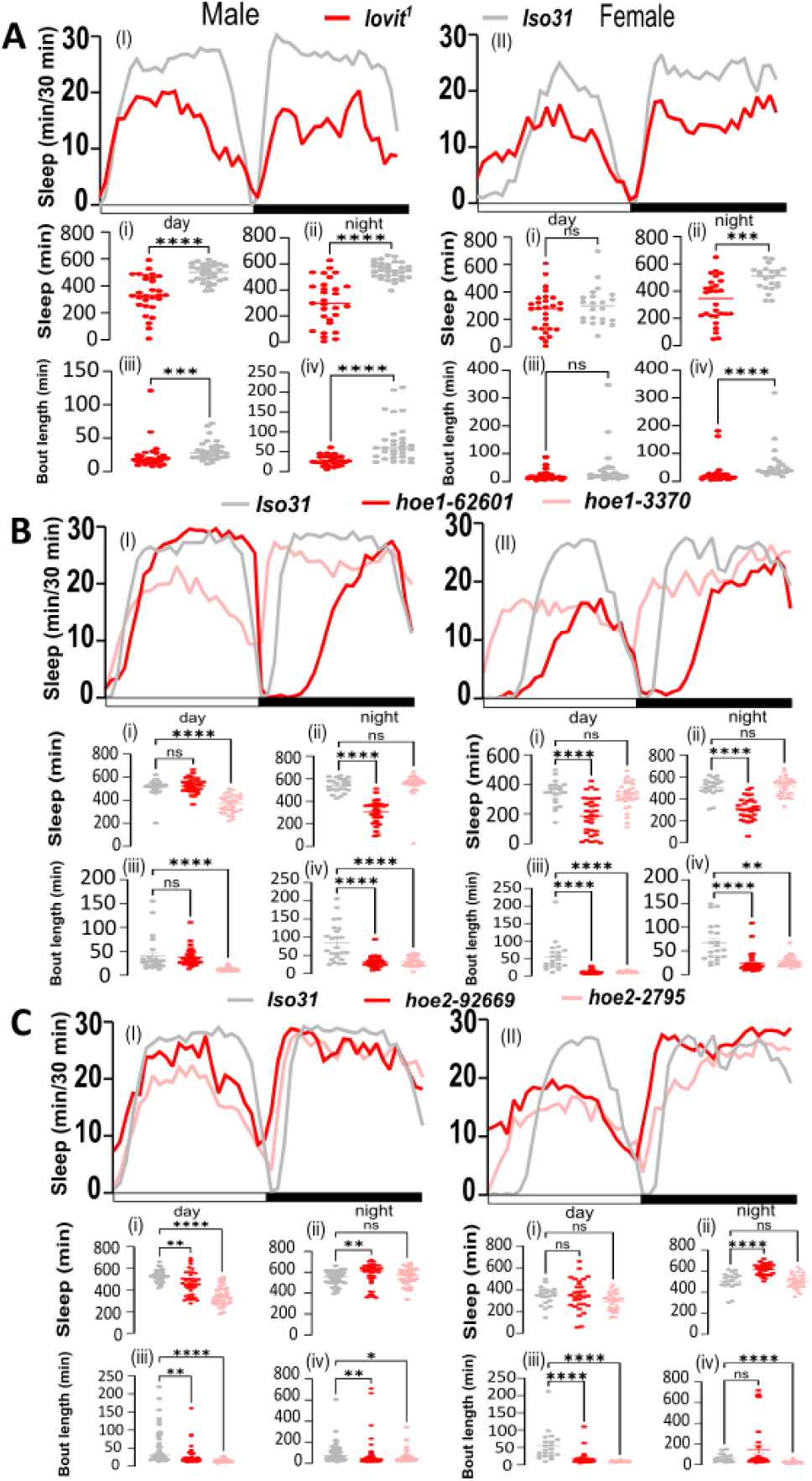
Sleep parameters of *lovit^1^, hoe1* and *hoe2* mutants in flies. 24-hour sleep profile of flies expressing **(A)** *lovit^1^* and **(B)** *hoe1* mutants, and (**C**) *hoe2* mutants. The line graphs for male (**I**) and female (**II**) show sleep amount (minutes per 30-minute bin) across the LD cycle (0-720 light [day] and 720-1440 dark [night]). The scatter plots show total daytime sleep (**i**, minutes, 12-h light phase), total nighttime sleep (**ii,** minutes, 12-h dark phase), daytime average sleep bout length (**iii,** minutes), and nighttime average sleep bout length (**iv,** minutes). Data are presented as individual data points with mean ± SEM. Statistical comparisons were performed using the Mann-Whitney U test for *iso31 vs. lovit^1^* and Kruskal-Wallis test with Dunn’s multiple comparisons correction for *iso31 vs. hoe1/2*. ns, not significant; *P < 0.05; **P < 0.01; ***P < 0.001; ****P < 0.0001. ***lovit^1^ vs. iso31***: n = 31 and 28 for male and female *lovit^1^*. n = 31 and 22 for male and female *iso31*. ***iso31 vs. hoe1*:** n = 24 and 21 for male and female *iso31* and n = 31-32 per genotype for both *hoe1* mutants. ***iso31 vs. hoe2:*** n = 38 and 21 for male and female *iso31*, n = 47 and 30 for male and female *hoe2-92669*, and n = 31 for male and female *hoe2-2795*.

#### hoe1/2 and lovit mutant alleles reduce circadian rhythmicity

We further verified the quality of circadian locomotor rhythm for the five mutants by monitoring their spontaneous locomotor activity in constant darkness and assessing rhythmicity. We found that both male and female mutants showed significantly lower rhythmicity judged by the clear dampening of peak to trough amplitude (Figure 2A-B) and the significant reduction of the Rhythmic Statistics value (Figure 2C-D), the established autocorrelation based index for circadian rhythmicity (29). Moreover, a small lengthening of circadian lengths was detected from the rhythmic population of *hoe1/hoe2* and *lovit* male mutants (RS≥ 2) (0.5 - 2 hours, Figure 2E). Similar period lengthening was observed in female mutants except for *hoe1^3370^* and *hoe2^2795^*, likely due the reduced number of rhythmic flies available in these two lines.

**Figure 2.**
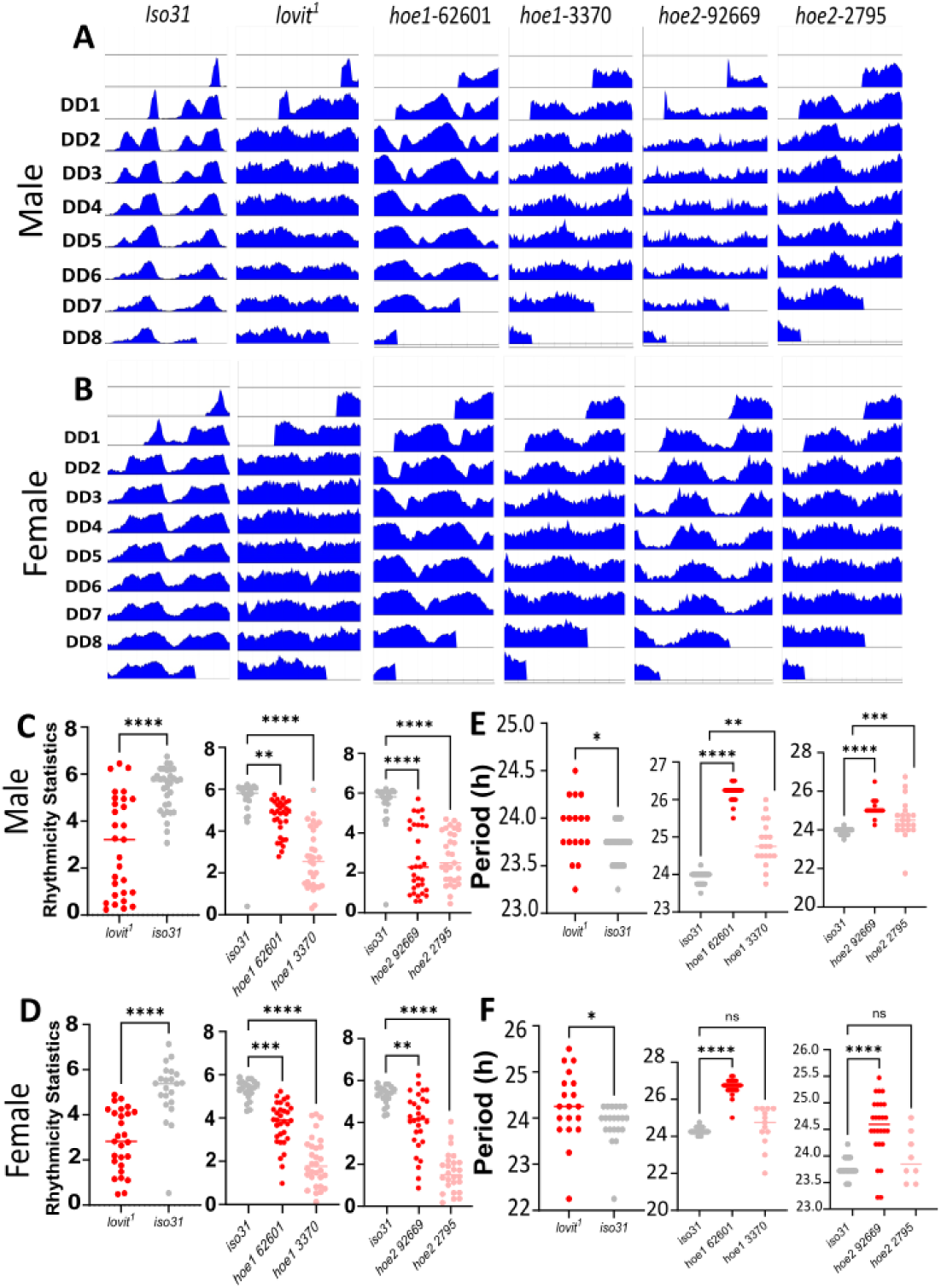
Circadian rhythm of locomotor activities for *lovit^1^, hoe1* and *hoe2* mutants. Double-plotted actograms for (**A**) male and (**B**) female flies showing eight consecutive days of activity in constant darkness (DD1–DD8) following from three days of 12:12 LD entrainment. Each panel shows the population-averaged activity across all flies in that condition. (**C-D**) Male and female RS values were calculated from seven days of DD activity recording (DD1–DD7) using the FlyToolBox in MATLAB. (**E-F**) Male and female period lengths were calculated for the rhythmic flies (RS >2) from seven days of DD activity recording (DD1–DD7) using the FlyToolBox in MATLAB. Each point represents one fly. Data are presented as individual data points with mean ± SEM. Statistical comparisons were performed using the Mann-Whitney U test for *iso31 vs. lovit^1^* and Kruskal-Wallis test with Dunn’s multiple comparisons correction for *iso31 vs. hoe1/2*. ns, not significant; *P < 0.05; **P < 0.01; ***P < 0.001; ****P < 0.0001. (**A-D**) ***lovit^1^ vs. iso31***: n = 29 for male *lovit^1^* and 27 for female *lovit^1^*. n = 31 for *iso31* male and 21 for *iso31* female. ***iso31 vs. hoe1*:** n = 24, 32, and 31 for male *iso31*, *hoe1-62601*, and *hoe1-3370*. n = 21, 32, and 30 for female *iso31*, *hoe1-62601*, and *hoe1-3370*. ***iso31 vs. hoe2:*** n = 24, 32, and 31 for male *iso31*, *hoe2-92669*, and *hoe2-2795*. n = 21, 29, and 26 for female *iso31*, *hoe2-92669*, and *hoe1-2795*. (**E-F**). ***lovit^1^ vs. iso31***: n = 17 for *lovit^1^* male and 19 for *lovit^1^* female. n = 32 for *iso31* male and 21 for *iso31* female. ***iso31 vs. hoe1*:** n = 23, 32, and 19 for male *iso31*, *hoe1-62601*, and *hoe1-3370*. n = 21, 30, and 13 for female *iso31*, *hoe1-62601*, and *hoe1-3370*. ***iso31 vs. hoe2:*** n = 23, 17, and 21 for male *iso31*, *hoe2-92669*, and *hoe2-2795*. n = 21, 26, and 8 for female *iso31*, *hoe2-92669*, and *hoe2-2795*.

### Electroretinograms reveal photoreceptor neurotransmission defects in *lovit¹* and *hoe1/2* **mutants**

The coincidental sleep and circadian phenotype observed in *hoe1/hoe2* and *lovit* mutants resemble previous findings in classic vision mutants (24). To test whether the sleep and circadian phenotypes observed reflect altered visual signalling, electroretinograms (ERGs) were recorded from *iso31*, *lovit*^1^, and four *hoe* mutant alleles (Figure 3). Typically, ON and OFF transient potential at the transition of light on/off reflect synaptic output of the photoreceptor while the large depression potential RP indicates the light induced depolarisation of the photoreceptor (Figure 3A).

**Figure 3.**
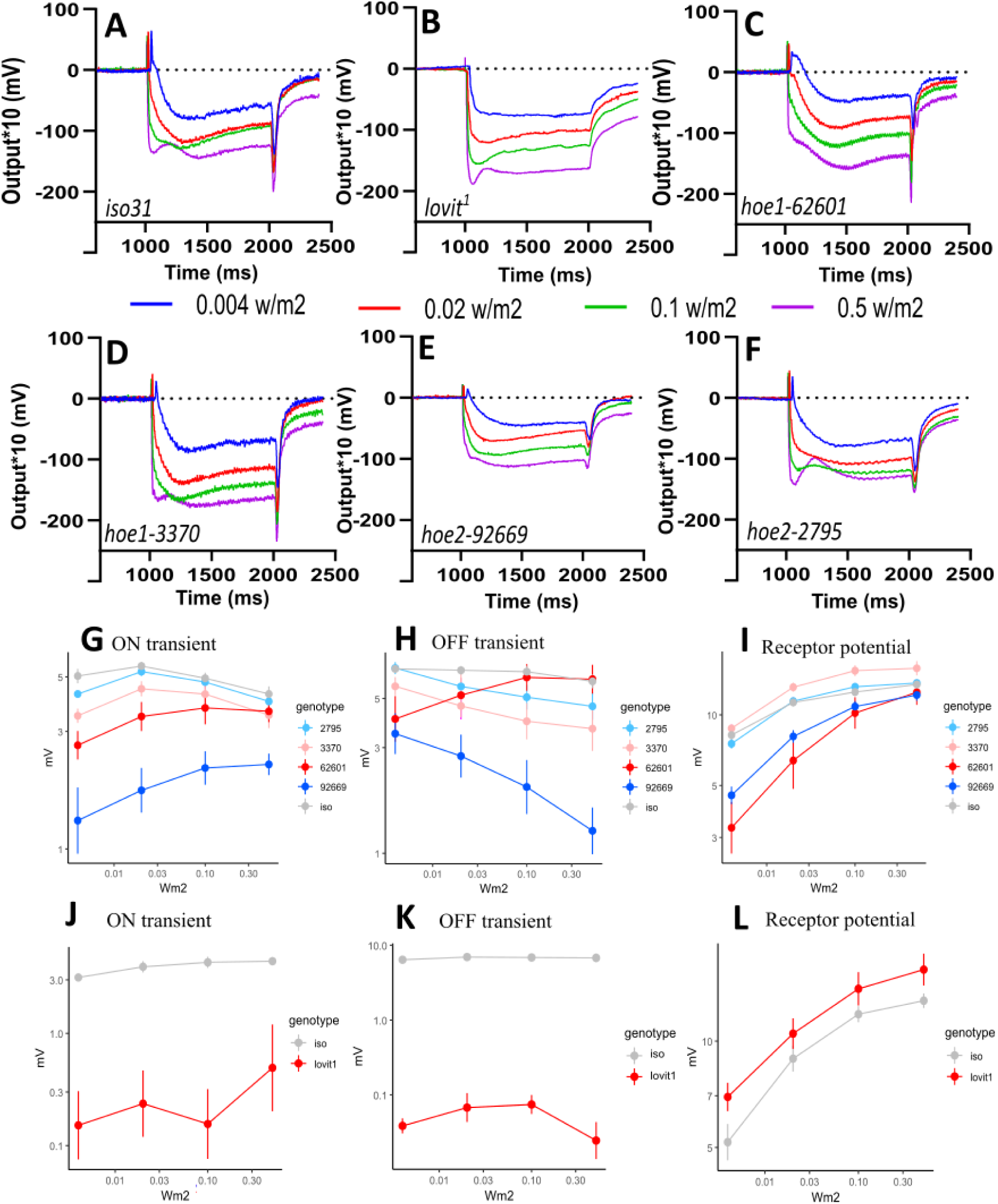
Electroretinogram assay for *lovit^1^, hoe1* and *hoe2* mutants. (**A-F**) Example ERG profiles of (**A**) *iso31*, (**B**) *lovit^1^*, (**C**) *hoe1-62601*, (**D**) *hoe1-3370*, (**E**) *hoe2-92669*, and (**F**) *hoe2-2795* with 1000 ms green light flash that evokes voltage changes of on-transient, receptor potential and off-transient upon four light intensities (0.004, 0.02, 0.1 and 0.5 w/m^2^). (**G-I).** Mean ± SEM for voltage change (mV) in on transient, receptor potential and off-transient of ERG upon four light intensities (0.004, 0.02, 0.1 and 0.5 w/m^2^) flash for *iso31* and *hoe1/2* mutants. (**J-L**) Mean ± SEM for voltage change (mV) in on transient, receptor potential and off-transient of ERG upon four light intensities (0.004, 0.02, 0.1 and 0.5 w/m^2^) flash for *lovit^1^* and *iso31*. Data for *iso31* in J-L are duplicated here from Figure 1I in (26) as the experiments were performed at the same time. The axis is plotted in logarithmic.

Consistent with previous findings (30), *lovit¹* ON and OFF transients were reduced as compared to control flies (see example traces, Figure 3B vs 3A, ON transient p = 1.6×10^−9^; ON r = 0.81, OFF transient p = 1.5×10^−6^; OFF r = 0.81, Table S4). Receptor potential was unchanged (p = 0.16, Figure 3J-L, Table S4). The selective loss of ON and OFF transients with preserved RP indicates a presynaptic neurotransmission defect, consistent with the known role of lovit protein as a vesicular histamine transporter at photoreceptor terminals (30).

Across the four *hoe* alleles, the strongest phenotypic changes were observed in flies carrying the *hoe2*^92669^ allele (Figure 3E): ON transient reduction (Figure 3G, p = 3.2×10^−10^; r = 0.84, Table S4) and OFF transient reduction (Figure 3H, p = 1.3×10^−9^; r = 0.83, Table S4). The 3370 *hoe1* deletion allele showed reduced ON transients (p = 4.2×10^−4^; r = 0.51) and reduced OFF transients (p = 3.8×10^−4^; r = 0.52) (Figure 3D, 3G and 3H). The *hoe1^62601^* PBac allele exhibited reduced ON transients (p = 1.2×10^−5^; r = 0.69, Table S4) but did not reach significance for OFF transients (p = 0.24, Table S4)(Figure 3C, 3G and 3H). The *hoe2^2795^*allele showed an OFF transient defect (p = 0.041; r = 0.33, Table 4) but no ON transient change (p = 0.11, Table 4) (Figure 3F, 3G and 3H). Receptor potential differences across *hoe* mutants were borderline and only *hoe2^92669^* showed reduced levels (Figure 3I, p=0.039; r=0.384, Table S4); the phenotype is principally at the level of synaptic transmission rather than phototransduction. Statistical results for ERG are summarised in Table S4.

In summary, the *hoe1* and *hoe2* mutants all showed defects in ON and/or OFF transients with preserved or near-preserved RP as seen in *lovit*^1^ mutants. This places *hoe1* and *hoe2* in the same functional category as *lovit*: genes required for photoreceptor neurotransmission rather than for photoreceptor depolarisation per se.

### Expression pattern of *hoe1/2* and *lovit* reveal the role in sleep regulation

Since *hoe1^62601^* and *hoe2^92669^* are both Gal4-enhancer traps, we therefore used them to drive UAS-myRFP to investigate the expression patterns of *hoe1* and *hoe2* (Figure 4). The *hoe1* promoter show clearly expression surrounding neuropiles and in the cortex area (magenta, red arrows, Figure 4A-B) in the brain areas close to lamina (La), medulla (Me), anterior optic tubercle(AoTu) and optical glomeruli (OpG). This expression pattern resembles that of glia subgroups in *Drosophila* brain (31). Consistently, *hoe1* expression is enriched in the four major subtypes of glia cells in the public scRNA-Seq data (32)(Figure 4E). Additionally neuronal expression of *hoe1* is detected in a small number of cells with large soma similar to neuropeptide expressing neurons (yellow triangles, Figure 4B). This is consistent with the increased expression of *hoe1* in CCHa-1 peptide expressing neurons (Figure 4E). The expression pattern of *hoe2* promoter is similarly detected in glia cells (red arrows, Figure 4C-D), and many sleep and vision-related areas including Mushroom Body (KC/MB), Fan shaped Body (FB), Eplisoid Body (EB), Pars Intercerebralis (PI) and Medulla (Me) (Figure 4C). In consistency with the promoter expression pattern, the scRNA-Seq analysis showed that *hoe2* is expressed in the visual system (including eye photoreceptors), glia, sleep/circadian centres and peptidergic neurons located in PI (Figure 4E). Moreover, *hoe2* expression is enriched in tyraminergic and octopaminergic cells, both of which are sleep regulators (33, 34). Intriguingly, our scRNA-Seq analysis also revealed that *lovit* expresses highly in tyraminergic, octopaminergic and dopamineric cells and Dm9 medulla neurons (Figure 4E), apart from its known photoreceptors expression (30). In summary, the expression pattern of *hoe1/2* and *lovit* support their role as sleep and vision regulators.

**Figure 4.**
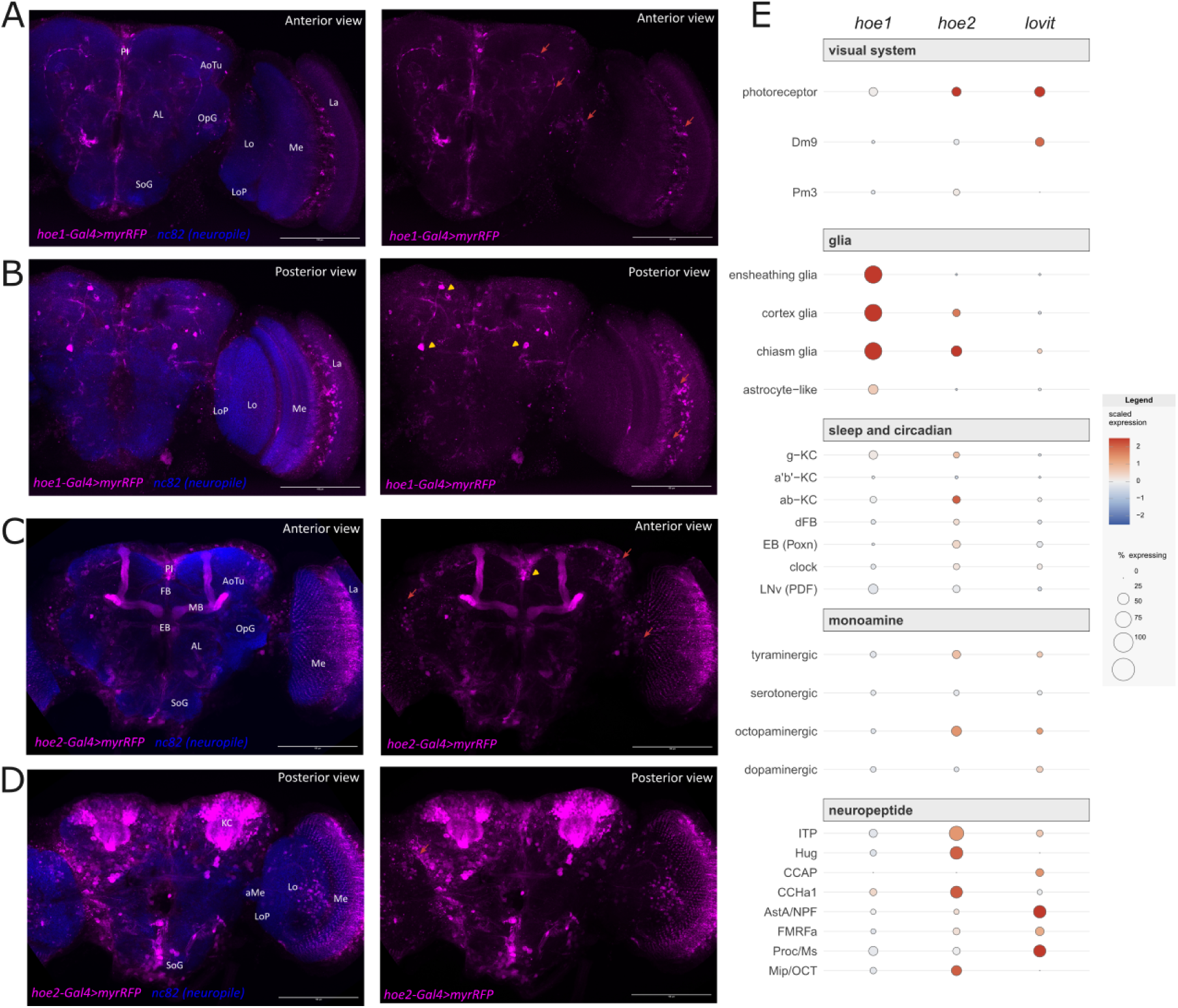
Spatial expression pattern of *hoe1*, *hoe2* and *lovit* in the fly brain. **A.** Anterior and **B.** Posterior views of *Drosophila* brain labeled by *hoe1-Gal4>myrRFP* (magenta) and nc82 (blue) showing *hoe1* promoter expressing cells and neuropiles**. C.** Anterior and **D.** Posterior views of *Drosophila* brain labeled by *hoe2-Gal4>myrRFP*(magenta) and nc82 (blue) showing *hoe2* promoter expressing cells and neuropiles. Red arrows indicate glial cell arborisations. Yellow triangles indicate neuropeptideric cells. The following neuropiles are indicated La (Lamina), Me (Medulla), Lo (Lobula), LoP (Lobula Plate), OpG, AoTu, MB (Mushroom Body)/KC (Keyon Cells), FB (Fan shaped Body), EB (Ellipsoid Body), AL (Antennal Lobes), SoG (suboesophageal ganglion), and PI (Pars Intercerebralis). Scale bar= 100um. **E.** Localisation of *hoe1*, *hoe2*, and *lovit* transcripts across curated *Drosophila* brain cell types. Dot plot showing detection and relative expression of the three candidate genes across published cell-type annotations (rows) from a single-nucleus RNA-seq atlas of the adult *Drosophila* brain (GSE107451), grouped into five functional categories: visual system, glia, sleep and circadian, monoamine, and neuropeptide. Dot size indicates the percentage of nuclei of that cell type in which the transcript was detected (≥1 count). Dot colour indicates scaled expression: for each gene, mean expression per cell type was z-scored (mean = 0, SD = 1, clipped at ±2.5) against all annotated cell types in the full dataset, not only those shown here, so colour values are directly comparable to the whole-dataset reference plot. Red indicates expression above the dataset-wide average for that gene; blue indicates below average; white indicates average expression.

## Discussion

Four findings emerge from this study. First, OCA2 and SLC45A2 orthologues, *hoe1/2* and *lovit,* regulate sleep in *Drosophila*. Second, germline mutations of *hoe1/2* and *lovit* do not change cuticle colouration, indicating that the sleep function of these genes is independent of melanin synthesis. Third, *hoe1/2* and *lovit* regulate photoreceptor neurotransmission as measured by ERG, linking the defects to the visual pathway. Fourth, *hoe1/2* and *lovit* express in vision or sleep regulatory brain areas.

### Pleiotropic sleep control by *OCA2* and *SLC45A2* orthologues

OCA2 and SLC45A2 orthologues are members of two broader transporter families which function in diverse cell types and subcellular locations (35, 36). Skin pigment cells and neurons share a common embryonic origin. In vertebrates, melanocytes derive from the neural crest, an ectodermal lineage that also gives rise to peripheral neurons (37). In invertebrates, cuticular epidermis cells and neurons both originate from the embryonic ectoderm. This shared developmental origin allows for pleiotropic functional diversification in which an ancestral transporter or channel function in pigmentation can be co-opted for neurons, or vice versa. Interestingly, based on human scRNA-seq and proteomic datasets (17, 18), human OCA2 and SLC45A2 (and potentially their vertebrate orthologues) are expressed in melanocytes as well as in the nervous system. Similarly, our imaging and scRNA-Seq analysis here places *hoe1* and *hoe2* in glial cells and neurons of the optic lobe and central brain, and *lovit* in the eye photoreceptors (30, 32) (Figure 4). The data showing that *hoe1/hoe2* and *lovit* mutants (null and RNAi knockdown) exhibit sleep but not pigmentation phenotypes are therefore consistent with the proposal that across evolution *Drosophila* retained the neuronal role of these transporters, while vertebrates maintain both skin pigmentation and neuronal function.

On the other hand, the lack of a sleep phenotype in RNAi-mediated knockdown for *CG31076*, *CG12061* and *Slc45-1* may be explained by evolutionary diversification in *Drosophila*. This may be the case as *Drosophila* lacks *TYR*, *TYRP1* and *DCT* orthologues and the loss of the *LRMDA* orthologue (*CG31076*) enhances rather than reduces pigmentation in *Drosophila*. Alternatively, it may be due to the cell type or developmental window not being targeted: the *ey*-Gal4 employed herein has limited expression outside the developing eye in *Drosophila*, and the tissue-specific expression patterns of CG31076 and CG12061 have not been characterized.

The day time sleep phenotypes observed in the *hoe1/hoe2* and *lovit* null mutants resembles previous findings that children with albinism scored lower on daytime sleepiness than other visually impaired children did (12), suggesting a light-sensing rather than image-forming mechanism may contribute to the OCA2/SLC45A2-mediated day sleep (see below). Our data are also consistent with the proposed pleiotropic sleep regulatory function of OCA2 orthologues in the *Astyanax* cavefish in which loss of *oca2* causes both albinism and sleep loss (15). These previous and our findings together imply a common OCA2/SLC45A2-mediated sleep regulatory mechanism conserved across deuterostomes and protostomes.

### Potential sleep regulatory pathway via ligand-gated chloride channels

LOVIT expression was previously characterised (30) and found mainly in the eye photoreceptors. Unlike SLC45A2, LOVIT protein acts as the vesicular histamine transporter that loads photoreceptor synaptic vesicles. Histamine is the main neurotransmitter in *Drosophila* photoreceptors and its synaptic transmission depends on the histamine-gated chloride channels (Ort/HisCl1) at postsynaptic neurons and glia cells. Therefore, loss of LOVIT reduces the ON and OFF transients of the ERG, leaving the RP intact. The data presented here reproduce this phenotype for *lovit*^1^ and importantly demonstrated that *hoe1* and *hoe2* mutants produce a phenotype of the same but weaker absolute magnitude or only affecting ON or OFF transients as compared to *lovit*^1^. As mentioned above, expressions of *hoe1* and *hoe2* are detected in the fly visual system and central brain (Figure 4). Particularly *hoe1* and *hoe2*-Gal4-mediated myrRFP reporters indicated that *hoe1* and *hoe2* express in optic chiasmic glial cells (Figure 4E). This group of glia wraps around photoreceptor synapses and maintains synaptic transmission partly through histamine recycling (38, 39). We have recently shown that reduction of photoreceptor outputs either by mutating phototransduction components or synaptic output causes sleep loss or sleep fragmentation in *Drosophila* (26). Considering the known OCA2 function as a proton-chloride co-transporter (5) and the coincidence between sleep and ERG phenotypes observed in *hoe1/2* and *lovit* mutant, our data therefore suggest that *hoe1* and *hoe2* may contribute to the ionic environment required for the histamine synthesis and recycling pathway (38–41). An alternative is that *hoe1*/2 support synaptic chloride homeostasis in the lamina neurons and in the direct postsynaptic target of eye photoreceptors. While histamine is a well-characterised arousal signal in mammals, SLC45A2 was demonstrated a proton-sucrose coupled symporter (6), but whether the same pH regulatory role persists in the mammalian neurons or it instead also acts as synaptic vesicular histamine transporter remains to be tested. Nevertheless, histamine mainly acts on postsynaptic G-protein couple receptors in mammals, and only at a higher concentration, histamine cross-talk to activate other ligand-gated channels (42). These channels include GABA_A_-gated chloride channels, whose neuronal inhibitory effects depend upon intracellular chloride levels, and are important for sleep and circadian regulation (43, 44). Therefore, the vertebrate OCA2 orthologues may control sleep via GABAergic pathways. Intriguingly, *hoe2* expresses in various sleep regulatory areas that required GABA_A_ signalling, including clock neurons(44), MB(45), FB(46) and EB(47) (Figure 4C-E). Future investigation of *hoe2* in these neurons will clarify if GABAergic signal is a conserved pathway for OCA2-mediated sleep.

As mentioned above the neuromelanin pathway and *Drosophila* cuticle melanin synthesis use similar dopamine/DOPA-dependent mechanisms. Indeed, a recent RNAi-based screen in the fruit fly identified a set of cuticle pigmentation regulators that affect dopamine and sleep (48). Although *Drosophila* OCA2 and SLC45A2 orthologues do not control cuticle pigmentation, their expressions are significantly detected in monoamineric and peptidergic neurons that are related for sleep regulation (Figure 4E). The fact that lovit protein can function as alternative Vesicular Monoamine Transporter (VMAT) for histamine, suggests a future investigation of the possibility that hoe1/2 and lovit may regulate sleep via other monoamines or neuropeptides in both synaptic vesicle and dense-core vesicles transmission within designated neural domain identities (49).

### *hoe1/2* and *lovit* control sleep and circadian rhythm via light or visual inputs

*Drosophila* sleep in both the day and night, with our previous findings suggesting that photoreceptors promote daytime sleep through the R1–R6 eye photoreceptors and the L2 lamina pathway (26). This is consistent with the current finding that the majority of *hoe1/2* and *lovit* null mutants (with the exception of *hoe1* 62601 males and *lovit^1^* females) have daytime sleep defects. Notably, the 62601 *hoe1* allele has a PiggyBac insertion in a non-coding intron and a phenotype that is weaker on ERG OFF transients than the 3370 deletion alleles. Further clarification of this allele as loss-of-function will be required. Similarly, our *ey*-Gal4 driven RNAi knockdown data are less consistent. This could be due to the RNAi efficiency as well as the developmental window and cell specificity (*ey*-Gal4 expression stops shortly after neuron and glial differentiation (50)). Nevertheless, the consistent day sleep defect from two independent *hoe1/2* alleles from different genetic backgrounds validated the role of *hoe1/2* in the day sleep regulation. The night sleep phenotype is less clear and may depend on complicated downstream circuital defects (26).

Consistent with the role of light as a signal to synchronise and maintain circadian rhythmicity in *Drosophila* (24, 51), we found that *hoe1/2* and *lovit* null mutants showed clear reduction of circadian rhythmicity. Further investigation will be required to clarify if the sleep loss observed depends on this circadian rhythmicity defect.

Nevertheless, the loss of photoreceptor neurotransmission as seen in *hoe1/2* and *lovit* mutants, irrespective of mechanism, would degrade the light input that is required to drive sleep and circadian rhythmicity. This places the *lovit*, *hoe1* and *hoe2* sleep phenotype in a mechanistically coherent framework: light input mediated sleep.

The concurrent visual transmission defects and sleep changes in *hoe1/2* and l*ovit* suggest that altered image forming visual input as well as light perception can underlie the sleep phenotype observed since visual experience drives sleep need in *Drosophila* and vertebrates (22, 25, 52). Further experimentation integrating visual load, either via circuital or visual environmental manipulation (25), will be required to test this possibility.

## Conclusion

hoe1/2 and lovit regulates sleep, circadian rhythm and photoreceptor neurotransmission in *Drosophila*. The majority of *hoe1/2* and *lovit* knockout alleles showed sleep defects and altered circadian rhythmicity in both sexes. ERG recordings show that *hoe1/2* and *lovit* alleles share a presynaptic neurotransmission defect phenotype with reduced ON and/or OFF transients and preserved receptor potential. The lack of pigmentation phenotype suggests OCA2 and SLC45A2 orthologues in *Drosophila* retain pleiotropic neuronal and sleep regulatory function, which may be related to the sleep symptoms documented in patients with oculocutaneous albinism.

## Materials and Methods

### Orthologue identification

Human OCA-associated genes for OCA1–8 were identified from the Online Mendelian Inheritance in Man database (OMIM). The DRSC Integrative Ortholog Prediction Tool version 10.0 (DIOPT) (53, 54) was used to identify *Drosophila melanogaster* orthologues. DIOPT integrates predictions from 13 databases and reports candidates with bidirectional best-hit (“yes-yes”) calls when the candidate is the highest-ranked orthologue in both human-to-fly and fly-to-human directions. Orthologues meeting the bidirectional criterion were selected for screening.

### *Drosophila* stocks and tissue specific knockdown of OCA orthologues

Fly food and husbandry were as previously described (26). Stocks were maintained on standard food at 25 °C under a 12 h:12 h light:dark cycle. Long-term stocks were kept on the same food at 18 °C. Prior to the sleep experiment, strains and crosses were transferred to a richer rearing food for better fly health. All fly lines, except for the *iso31* control (26), were purchased from stock centres (BDSC, VDRC and NIG, Table S1). The *lovit*^1^ allele was further outcrossed to *iso31* by swapping the X/Y and second chromosomes with those of *iso31*. The *hoe1^62601^* allele carries a PiggyBac insertion in a non-coding intron and may not abolish gene function; the *hoe2^92669^* allele is a CRIMIC insertion that disrupts the coding sequence. the *hoe1^3370^* and *hoe2^22795^* are CRISPR-mediated loss-of-function alleles.

The Gal4-UAS system was used to reduce OCA orthologues in developing eyes (55). The *eyeless* promoter-fused Gal4 (*ey-Gal4*, Table S1) expresses in the primordium eye tissue in the developing eye-antennal imaginal disc (56). Virgin *ey*-Gal4 females were crossed to UAS-RNAi males, and F1 progeny with correct genotype were assayed.

### Sleep and circadian assays

Sleep recordings were performed as described in Hung et al (26). Three- to four-day-old mated males and mated females were transferred individually to behaviour tubes (2% w/v agar, 4% w/v sucrose at one end, cotton wool at the other) and loaded into *Drosophila* Activity Monitors (DAM2, TriKinetics Inc., Waltham, MA, USA). Monitors were housed in a temperature- and light-controlled incubator (MIR-254-PE, PHCbi) at 25°C under a 12 h:12 h light:dark (LD) cycle for three days. Sleep was defined as continuous immobility ≥ 5 minutes. Day 3 of recording was used for sleep analysis to ensure the flies are climatised and circadian entrained to the condition. Both sexes were recorded. Beam-crossing data from the DAM monitors were processed using the customised Excel calculators (26). The calculator outputs five primary measures: sleep profile in 30-minute bins across 24 hours, daytime sleep (minutes of sleep per 720 minutes of light phase), nighttime sleep (minutes of sleep per 720 minutes of dark phase), daytime sleep bout length, and nighttime sleep bout length. Processed data were plotted in GraphPad Prism v10 (GraphPad Software, San Diego, CA, USA). Circadian locomotor activity was recorded for 7 to 8 days in constant darkness following the 3 days of LD cycles. Rhythmicity was analysed as previously described (29) by the autocorrelation-based parameter, Rhythmic Statistic (R.S.) for each genotype, while the actual period was scored based on the time interval at the second significant correlation coefficient peak.

### Electroretinogram recording and analysis

Electroretinograms (ERG) were recorded as previously described (26) with modification. Three- to five-day-old, mated male flies were used: *iso31*, *lovit*^1^ (*iso31* background), *hoe1* alleles *3370* and *62601*, and *hoe2* alleles *2795* and *92669*. *iso31* served as the genetic background control, and *lovit*^1^ served as a positive control for visual neurotransmission loss (30). Three components were quantified from the trace: the ON transient and OFF transient (representing photoreceptor-to-lamina synaptic responses) and the receptor potential (RP, the sustained depolarisation of the photoreceptor cell body during light exposure). A defect restricted to ON and OFF transients with preserved RP indicates a presynaptic neurotransmission defect rather than a phototransduction defect.

### Confocal imaging

Adult male flies (3–5 days old) were anesthetized at 4°C, and brains were dissected in cold 1× PBS before immediate transfer to ice-cold 4% paraformaldehyde in PBST (PBS containing 0.3% Triton X-100). One set of dissection was completed within 20 min, followed by fixation in fresh 4% paraformaldehyde/PBST for 40 min at room temperature. Brains were rinsed once and washed in PBST for 30 min before blocking in 5% goat serum/PBST. Samples were incubated with primary antibodies with 5% goat serum/PBST overnight at 4°C, washed three times with PBST, and incubated with Alexa Fluor-conjugated secondary antibodies in 5% goat serum/PBST overnight at 4°C. After three additional PBST washes, brains were mounted in VECTASHIELD Antifade Mounting Medium (H-1000) on microscope slides, stored at −20°C, and imaged using a Zeiss LSM 980 inverted confocal microscope. Primary antibody concentrations were as follows: mouse anti-BRP (nc82, Developmental Studies Hybridoma Bank) - 1:50; rabbit anti-DsRed Polyclonal Antibody (Takara) - 1:200. Alexa Fluor secondaries (Invitrogen) were used as follows: Alexa Fluor 647 goat anti-mouse IgG - 1:200 and Alexa Fluor 555 goat anti-rabbit IgG - 1:200.

### sc-RNA-seq analysis

#### Preprocessing

Single-cell RNA-seq data from the adult *Drosophila melanogaster* whole-brain atlas ((32)GEO: GSE107451) were analysed in R (v4.5.3) using Seurat (v5.5.0). The raw expression matrix (10x Genomics MEX format) was imported with Read10X(), and author-provided cell-type annotations were merged onto cell barcodes with AddMetaData(); nuclei lacking a published annotation were excluded. Counts were log-normalised (NormalizeData(), scale factor = 1×10⁴). Gene symbols were mapped to the informal aliases used throughout the manuscript (e.g., CG45782 → *lovit*) for readability.

#### Candidate gene localisation

To profile *hoe1*, *hoe2*, and *lovit*, we curated a subset of 26 annotated cell types spanning five relevant functional groups: visual system (3 types), glia (4), sleep/circadian (7), monoaminergic (4), and peptidergic (8). Expression was visualised with Seurat’s DotPlot(), where dot size denotes the percentage of nuclei of a given cell type detecting the transcript, and dot colour denotes scaled expression — each gene’s mean expression per cell type, z-scored (clipped at ±2.5 SD) against **all annotated cell types in the full dataset** (32). Anchoring the z-score to the whole atlas, rather than recomputing it within each display subset, ensures that colour values are directly comparable across every panel (Figure 4E); a cell type’s colour therefore reflects its expression relative to the entire brain, not merely relative to its neighbours within a given panel.

### Statistical analysis

The sample sizes were based on convention in *Drosophila* sleep studies (see e.g.(26, 29)). For sleep data, pairwise comparisons between two genotypes were tested with the non-parametric Mann–Whitney U test. Comparisons among three or more genotypes were tested with the non-parametric Kruskal–Wallis test followed by post-hoc pairwise Dunn’s tests. These tests were performed in GraphPad Prism v10. For RNAi experiments, a phenotype was considered validated only if the experimental genotype differed significantly from both controls (Gal4/+ and UAS/+) at p < 0.05. ERG data were analysed as described in Hung et al.(26): Kruskal–Wallis with pairwise Wilcoxon tests and Benjamini–Hochberg adjustment, with effect sizes reported as r = Z/√N. No genotype blinding was done, but sleep experiments for null alleles (Figure 1) were independently validated by two lab members. The ERG experiments (Figure 3) were performed without knowledge of sleep phenotype.

## Acknowledgments

This study was funded by University of Leicester to M.A. (F100 PhD studentships), J.B. and F.H. (PGT and UG programme funding), an Magistère Européen de Génétique programme internship to N.M, a BBSRC New Investigator Scheme (BB/W014939/1) to K-F.C and Y-C. H. A and a Ulverscroft Foundation support to M.G.T.

## Author Contributions

K-F.C. – Conceptualization, Methodology, Investigation, Writing – Review & Editing, Visualization, Supervision, Resources, Project administration, Funding acquisition;

M.A. – Investigation, Visualization, Writing - Original Draft, Writing – Review & Editing;

M.G.T. – Conceptualization, Writing – Review & Editing, Resources, Supervision;

F.G. –Writing – Review & Editing, Resources, Supervision; Y-C.H. – Methodology, Investigation

C.L. – Methodology, Investigation

N.M., J.B., and F.H. – Investigation, Validation

