## supplementary information for "*Drosophila* orthologues of oculocutaneous albinism-associated genes regulate sleep and circadian rhythm via visual neurotransmission"

**This PDF file includes:**

Figures S1 to S6

Tables S1 to S4

**Figures**

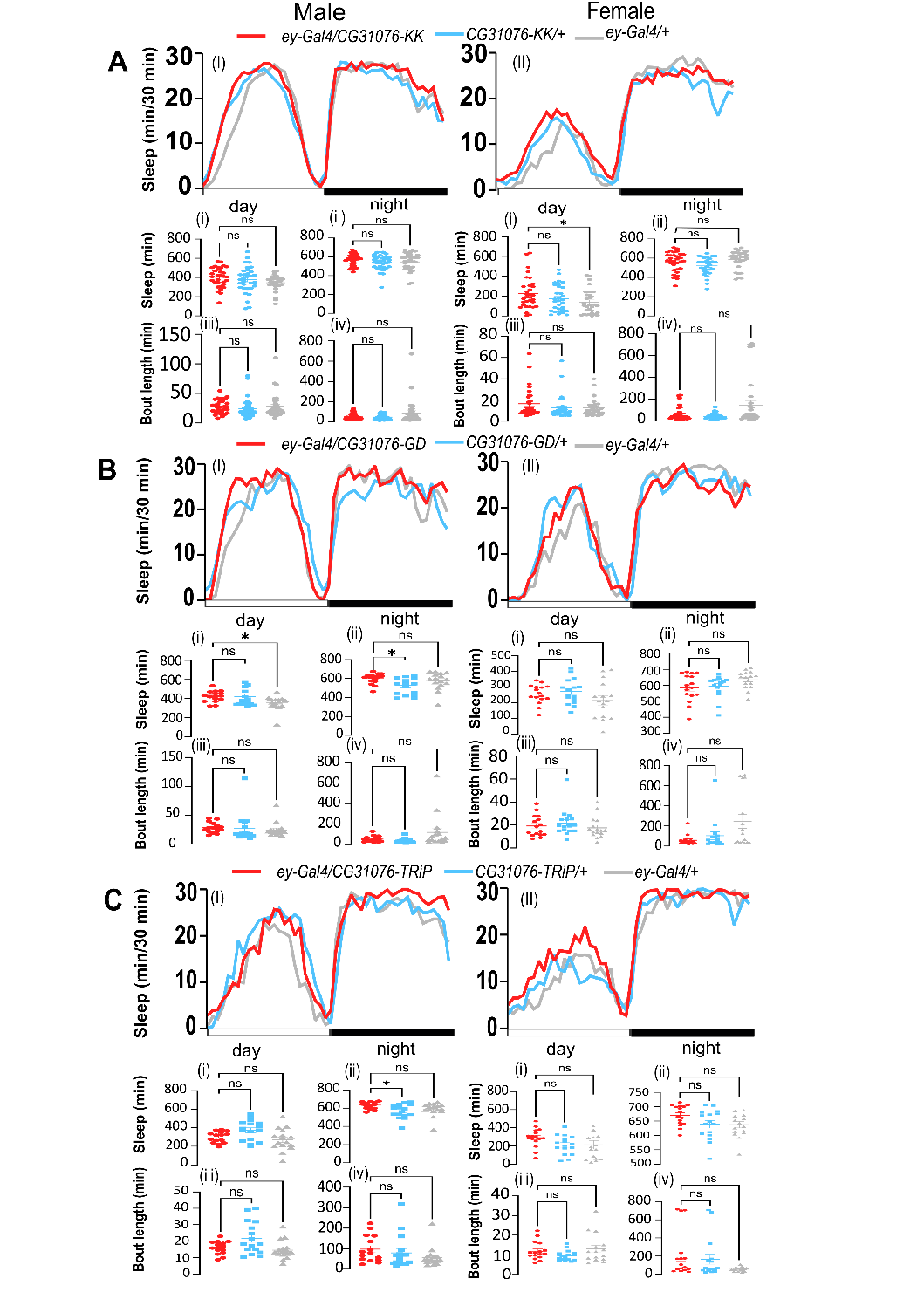

**Fig. S1.** ***ey-*Gal4-driven *CG31076* knockdown in flies.**

24-hour sleep profile of flies expressing **(A)** CG31076-KK, **(B)** CG31076-GD, and **(C)** CG31076-TRiP driven by *ey*-Gal4. The line graphs for male (**I**) and female (**II**) show sleep amount (minutes per 30-minute bin) across the LD cycle (0-720 light [day] and 720-1440 dark [night]). The scatter plots show total daytime sleep (**i**, minutes, 12-h light phase), total nighttime sleep (**ii,** minutes, 12-h dark phase), daytime average sleep bout length (**iii,** minutes), and nighttime average sleep bout length (**iv,** minutes).  Data are presented as individual data points with mean ± SEM. Statistical comparisons were performed using the Kruskal-Wallis test with Dunn’s multiple comparisons correction. ns, not significant; *P < 0.05; **P < 0.01; ***P < 0.001; ****P < 0.0001. n = 15-16 per genotype, except for CG31076-KK (n = 31-32 per genotype).

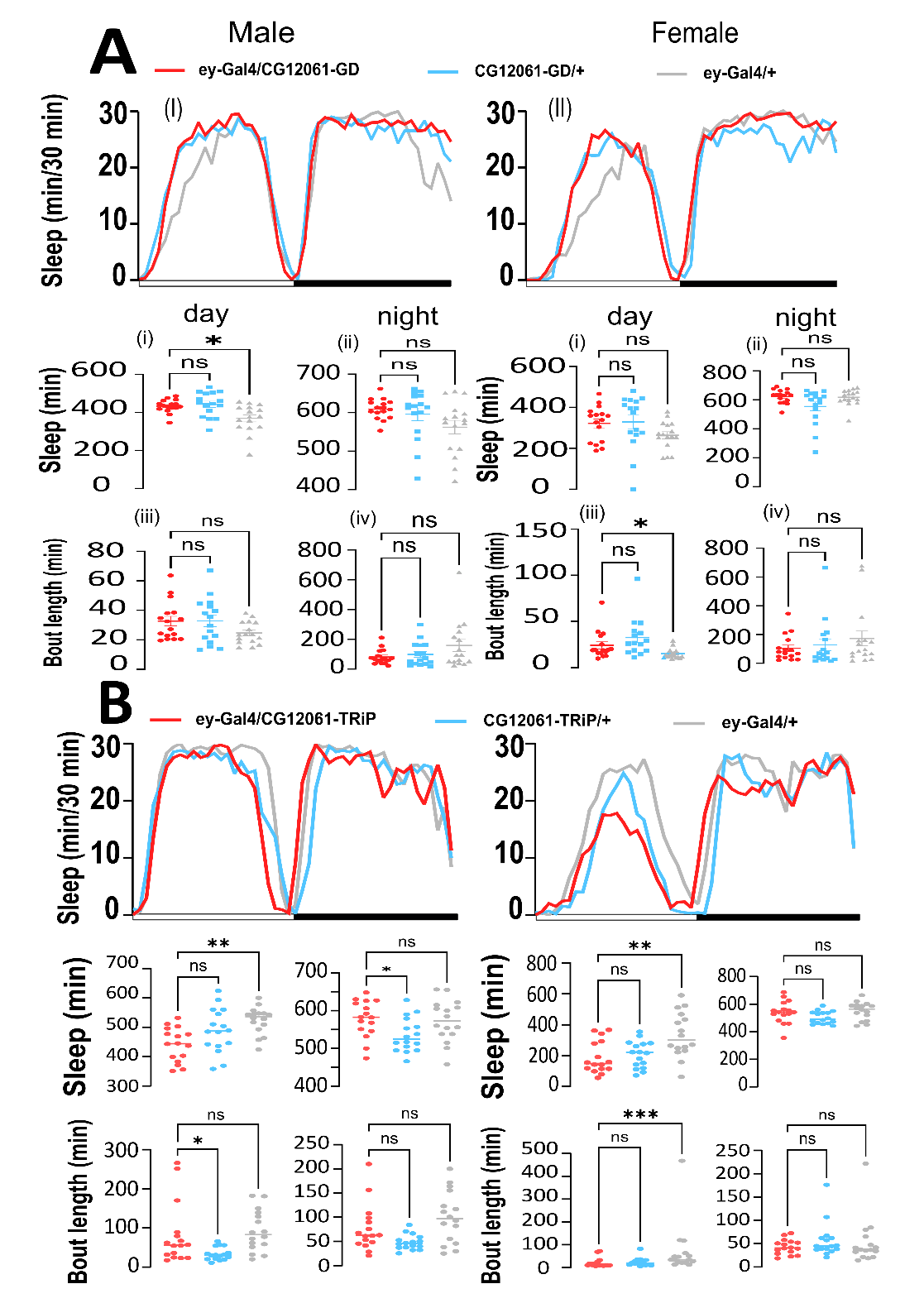

**Fig. S2.** ***ey-*Gal4-driven *CG12061* knockdown in flies.**

24-hour sleep profile of flies expressing **(A)** CG12061-GD **(B)** CG12061-TRiP driven by *ey*-Gal4. The line graphs for male (**I**) and female (**II**) show sleep amount (minutes per 30-minute bin) across the LD cycle (0-720 light [day] and 720-1440 dark [night]). The scatter plots show total daytime sleep (**i**, minutes, 12-h light phase), total nighttime sleep (**ii,** minutes, 12-h dark phase), daytime average sleep bout length (**iii,** minutes), and nighttime average sleep bout length (**iv,** minutes). Data are presented as individual data points with mean ± SEM. Statistical comparisons were performed using the Kruskal-Wallis test with Dunn’s multiple comparisons correction. ns, not significant; *P < 0.05. n = 16 per genotype.

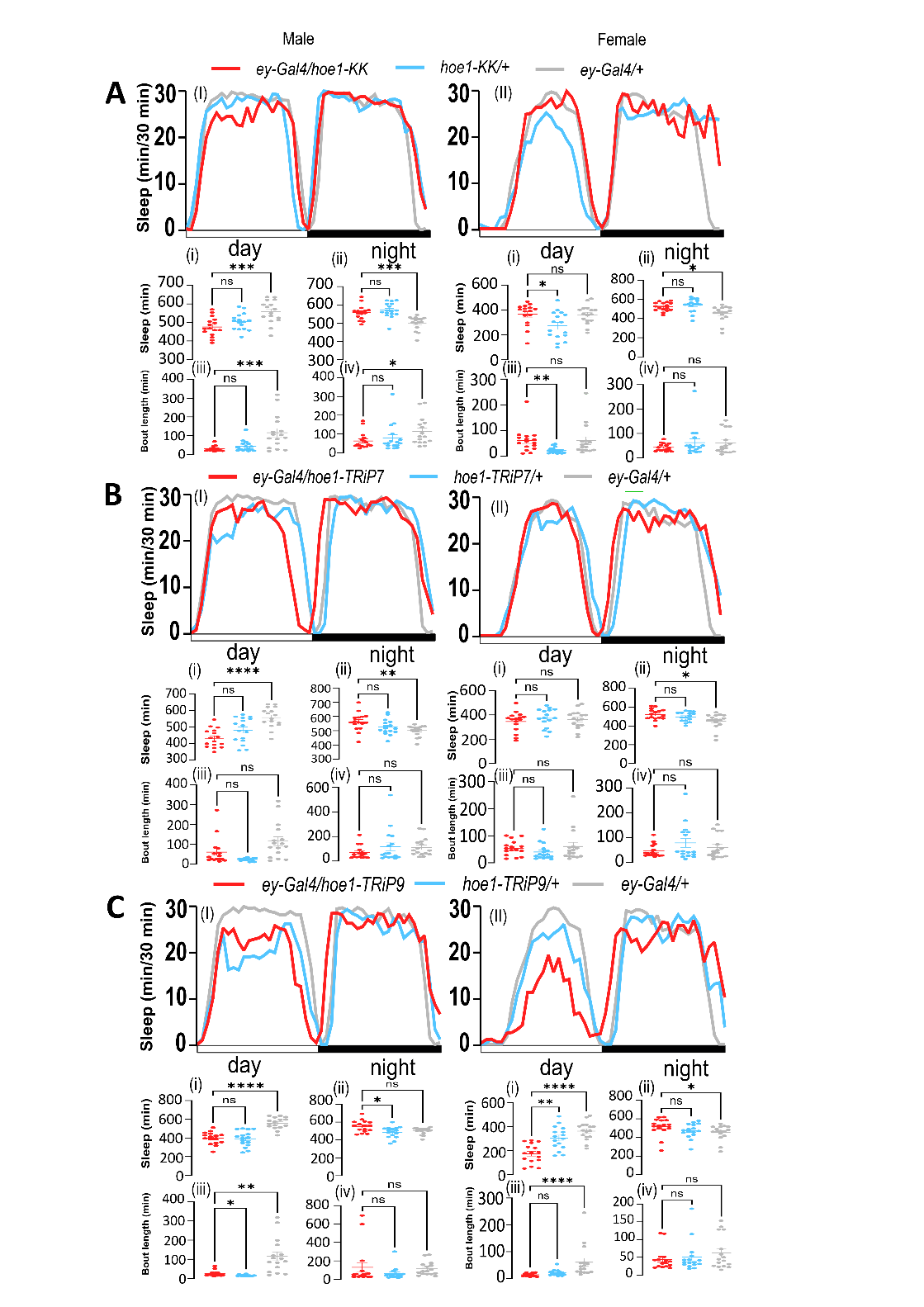

**Fig. S3.** ***ey-*Gal4-driven *hoe1* knockdown in flies.**

24-hour sleep profile of flies expressing **(A)** hoe1-KK, **(B)** hoe1-TRiP7, and **(C)** hoe1-TRiP9 driven by *ey*-Gal4. The line graphs for male (**I**) and female (**II**) show sleep amount (minutes per 30-minute bin) across the LD cycle (0-720 light [day] and 720-1440 dark [night]). The scatter plots show total daytime sleep (**i**, minutes, 12-h light phase), total nighttime sleep (**ii,** minutes, 12-h dark phase), daytime average sleep bout length (**iii,** minutes), and nighttime average sleep bout length (**iv,** minutes). Data are presented as individual data points with mean ± SEM. Statistical comparisons were performed using the Kruskal-Wallis test with Dunn’s multiple comparisons correction. ns, not significant; *P < 0.05; **P < 0.01; ***P < 0.001; ****P < 0.0001. n = 16 per genotype.

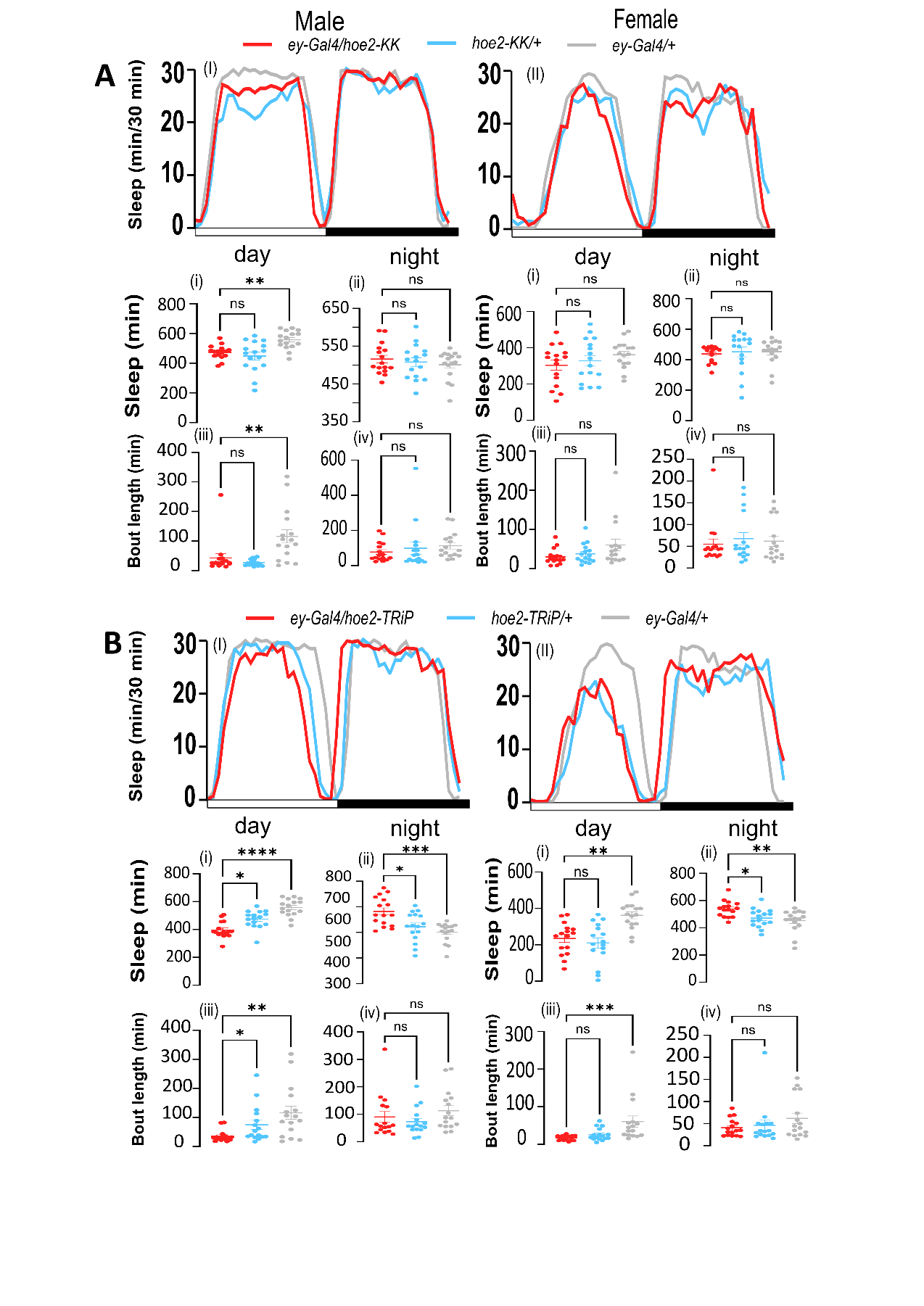

**Fig. S4.** ***ey-*Gal4-driven *hoe2* knockdown in flies.**

24-hour sleep profile of flies expressing **(A)** hoe2-KK and **(B)** hoe2-TRiP driven by *ey*-Gal4. The line graphs for male (**I**) and female (**II**) show sleep amount (minutes per 30-minute bin) across the LD cycle (0-720 light [day] and 720-1440 dark [night]). The scatter plots show total daytime sleep (**i**, minutes, 12-h light phase), total nighttime sleep (**ii,** minutes, 12-h dark phase), daytime average sleep bout length (**iii,** minutes), and nighttime average sleep bout length (**iv,** minutes). Data are presented as individual data points with mean ± SEM. Statistical comparisons were performed using the Kruskal-Wallis test with Dunn’s multiple comparisons correction. ns, not significant; *P < 0.05; **P < 0.01; ***P < 0.001; ****P < 0.0001. n = 16 per genotype.

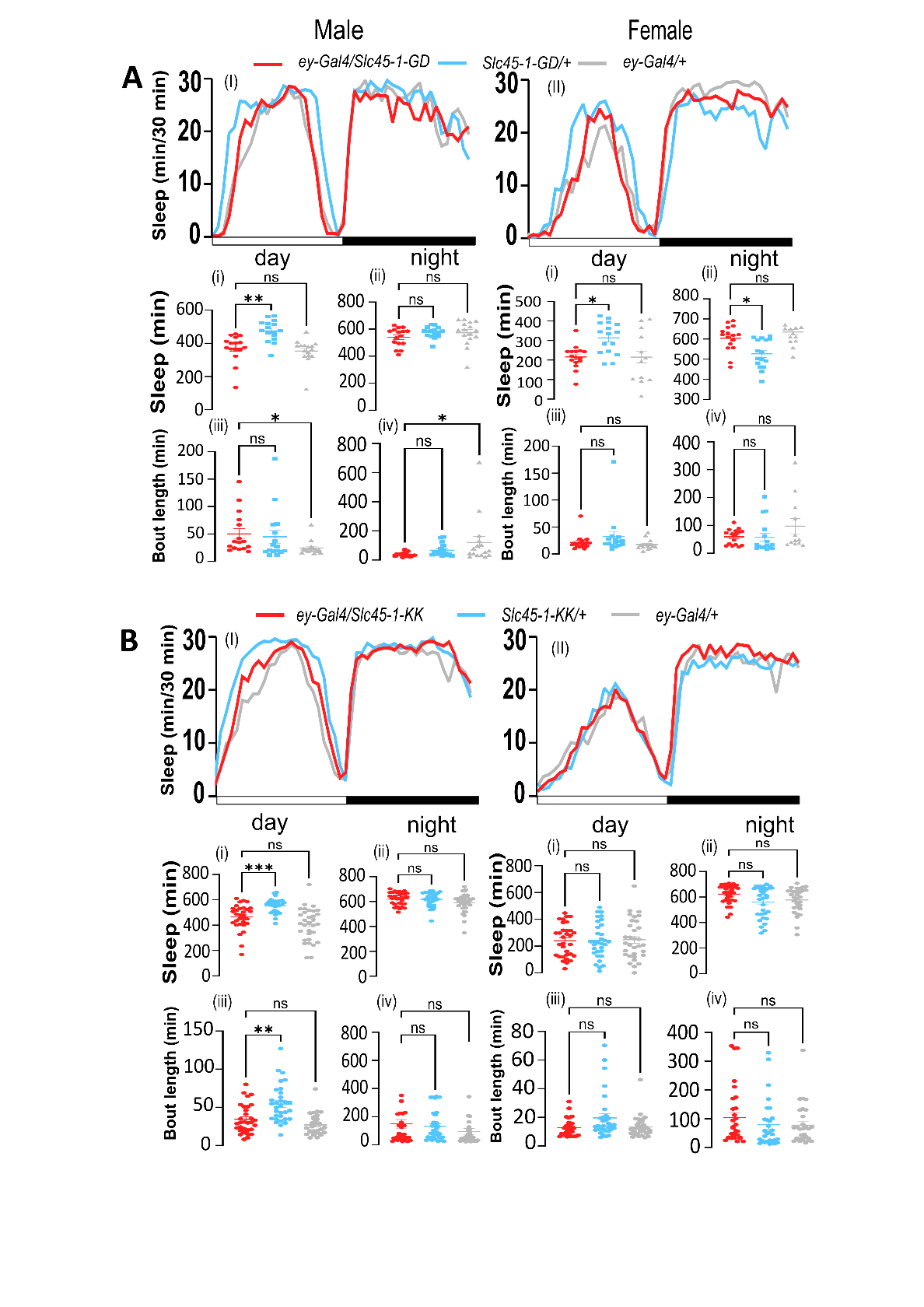

**Fig. S5.** ***ey-*Gal4-driven *Slc45-1* knockdown in flies.**

24-hour sleep profile of flies expressing **(A)** Slc45-1-GD and **(B)** Slc45-1-KK driven by *ey*-Gal4. The line graphs for male (**I**) and female (**II**) show sleep amount (minutes per 30-minute bin) across the LD cycle (0-720 light [day] and 720-1440 dark [night]). The scatter plots show total daytime sleep (**i**, minutes, 12-h light phase), total nighttime sleep (**ii,** minutes, 12-h dark phase), daytime average sleep bout length (**iii,** minutes), and nighttime average sleep bout length (**iv,** minutes). Data are presented as individual data points with mean ± SEM. Statistical comparisons were performed using the Kruskal-Wallis test with Dunn’s multiple comparisons correction. ns, not significant; *P < 0.05; **P < 0.01; ***P < 0.001; ****P < 0.0001. n = 16 for Slc45-1-GD (except for female *ey-Gal4/+*, for which n = 12) and 29-32 for Slc45-KK per genotype.

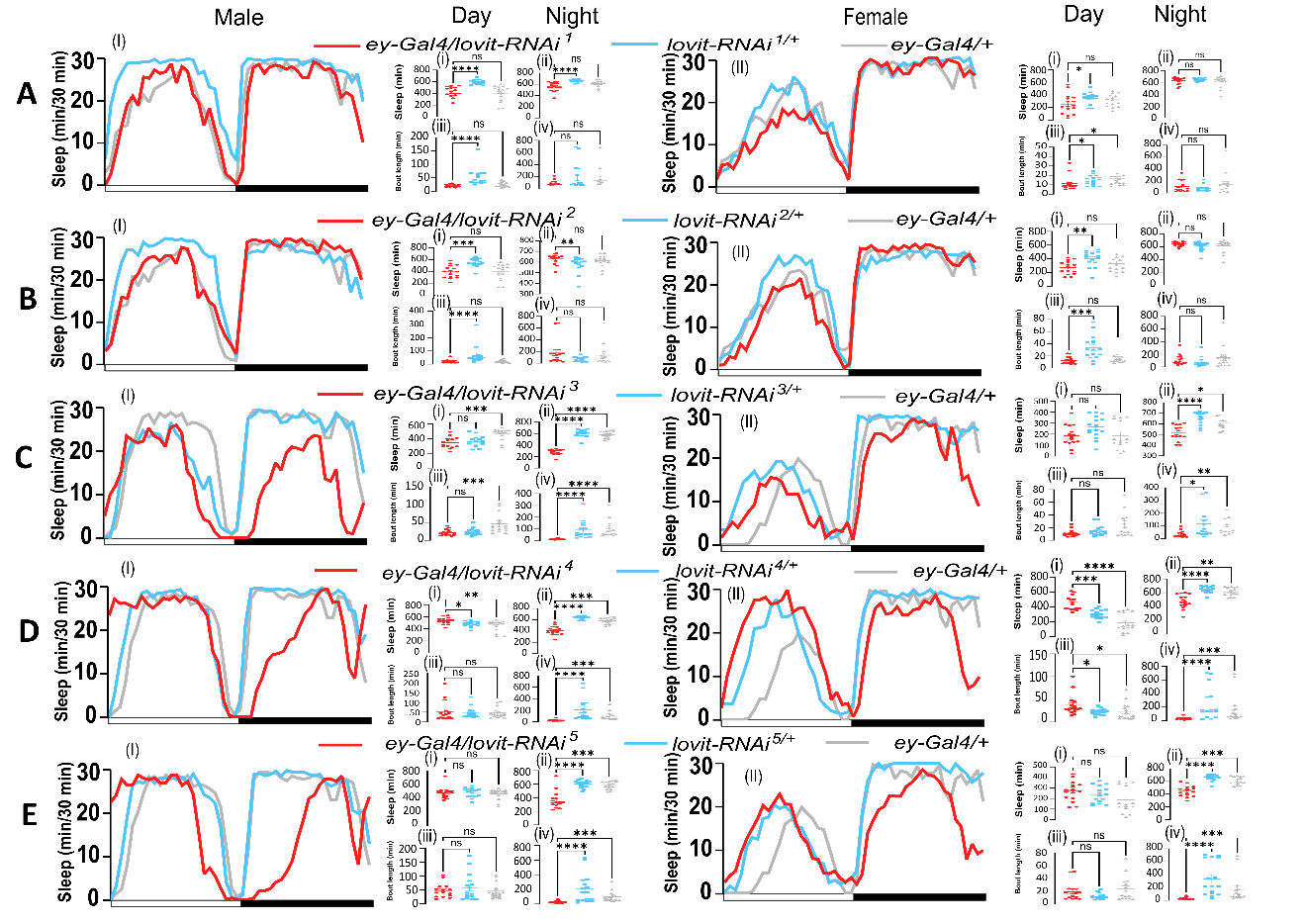

**Fig.S6.** ***ey-*Gal4-driven *lovit* knockdown in flies.**

24-hour sleep profile of male and female flies expressing **(A)** lovit-RNAi^1^, **(B)** lovit-RNAi^2^, (**C**) lovit-RNAi^3^, (**D**) lovit-RNAi^4^, and (**E**) lovit-RNAi^5^ driven by *ey*-Gal4. The line graphs for male (**I**) and female (**II**) show sleep amount (minutes per 30-minute bin) across the LD cycle (0-720 light [day] and 720-1440 dark [night]). The scatter plots show total daytime sleep (**i**, minutes, 12-h light phase), total nighttime sleep (**ii,** minutes, 12-h dark phase), daytime average sleep bout length (**iii,** minutes), and nighttime average sleep bout length (**iv,** minutes).  Data are presented as individual data points with mean ± SEM. Statistical comparisons were performed using the Kruskal-Wallis test with Dunn’s multiple comparisons correction. ns, not significant; *P < 0.05; **P < 0.01; ***P < 0.001; ****P < 0.0001. n = 15-16 per genotype.

**Tables**

**Table S1. *Drosophila* lines used in this study.**

Manuscript Name	Source ID (Stock center)	Target Gene

*iso31* control	See [26]	isogenised control

*CG12061-GD*	v8681 (VDRC)	*CG12061*

*CG12061-TRiP*	26248 (BDSC)	*CG12061*

*CG31076-TRiP*	60478 (BDSC)	*CG31076*

*CG31076-GD*	v28776 (VDRC)	*CG31076*

*CG31076-KK*	v107532 (VDRC)	*CG31076*

*lovit^1^*	83374 (BDSC)	*lovit*

*lovit-RNAi1*	v109094 (VDRC)	*lovit*

*lovit-RNAi2*	v109249 (VDRC)	*lovit*

*lovit-RNAi3*	38367 (BDSC)	*lovit*

*lovit-RNAi4*	40876 (BDSC)	*lovit*

*lovit-RNAi5*	61340 (BDSC)	*lovit*

*Slc45-1-GD*	v5174 (VDRC)	*Slc45-1*

*Slc45-1-KK*	v105439 (VDRC)	*Slc45-1*

*hoe1-KK*	v110373 (VDRC)	*hoe1*

*hoe1-TRiP7*	33377 (BDSC)	*hoe1*

*hoe1-TRiP9*	31719 (BDSC)	*hoe1*

*hoe1^62601^*	62601 (BDSC)	*hoe1*

*hoe1^3370^*	M2L-3370 (NIG)	*hoe1*

*hoe2-KK*	v106723 (VDRC)	*hoe2*

*hoe2-TRiP*	28661 (BDSC)	*hoe2*

*hoe2^92669^*	92669 (BDSC)	*hoe2*

*hoe2^2795^*	M2L-2795 (NIG)	*hoe2*

*uas-myrRFP*	7118 (BDSC)	*myrRFP*

*ey-Gal4*	5534 (BDSC)	Gal4-driver

**Table S2. Summary of RNAi screen sleep phenotypes.**

**Genotype	Lines	Day sleep	Night sleep	Day bout	Night bout**

*ey>CG31076 RNAi (LRMDA)*	3	n.s.	n.s.	n.s.	n.s.

*ey>CG12061 RNAi (SLC24A5)*	2	n.s.	n.s.	n.s.	n.s.

*ey>hoe1 RNAi*	3	Y (1/3)	n.s.	Y (1/3)	n.s.

*ey>hoe2 RNAi*	2	Y (1/2)	n.s.	Y (1/2)	n.s.

*ey>Slc45-1 RNAi*	2	n.s.	n.s.	n.s.	n.s.

*ey>lovit RNAi*	5	n.s.	Y (3/5)	n.s.	Y (3/5)

“Y” indicates significance against both controls; “n.s.” indicates not significant against both controls; “(1/2, 1/3, etc.)” indicates significance in one of two RNAis, one of three RNAis, etc.

**Table S3. Sleep parameters for hoe1/2 and lovit knockout alleles.**

**Genotype	Sex	Day sleep	Night sleep	Day bout	Night bout**

hoe1^3370^	M	↓	—	↓	↓

hoe1^62601^	M	—	↓	—	↓

hoe2^2795^	M	↓	—	↓	↓

hoe2^92669^	M	↓	↑	↓	↓

lovit¹	M	↓	↓	↓	↓

hoe1^3370^	F	—	—	↓	↓

hoe1^62601^	F	↓	↓	↓	↓

hoe2^2795^	F	—	—	↓	↓

hoe2^92669^	F	—	↑	↓	—

lovit¹	F	—	↓	—	↓

Direction of significant change vs. *iso31* control. ↓ reduced; ↑ increased; — not significant.

**Table S4. Electroretinogram results vs. iso31 control.**

**Genotype (n)	ON transient p	ON r	OFF transient p	OFF r	RP p	RP r	Magnitude (ON/OFF)**

*lovit^1^* (4)	1.6×10^-9^	0.81	1.5×10^-6^	0.81	0.16 (n.s.)	N.A.	Large /Large

*hoe1^3370^* (6)	4.2×10^-4^	0.51	3.8×10^-4^	0.52	0.053 (n.s.)	N.A.	Large /Large

*hoe1^62601^* (5)	1.2×10^-5^	0.69	0.243 (n.s.)	0.19	0.055 (n.s.)	N.A.	Large /N.A.

*hoe2^2795^* (6)	0.112 (n.s.)	0.23	0.041	0.33	0.84 (n.s.)	N.A.	N.A./Moderate

*hoe2^92669^* (4)	3.2×10^-10^	0.84	1.3×10^-9^	0.83	0.039	0.384 (moderate)	Large/Large

p-values from pairwise Wilcoxon test with Benjamini–Hochberg adjustment; effect size r reported as Z/√N. *lovit¹* vs. *iso31*: Kruskal–Wallis N.A. (two groups). *hoe1/2* alleles vs. *iso31*: Kruskal–Wallis ON p = 8.3×10^-12^; OFF p = 3.6×10^-8^; RP p = 3.1×10^-4^. N.A.: not available
